# Data-intensive modeling of forest dynamics

**DOI:** 10.1101/005009

**Authors:** Jean F. Liénard, Dominique Gravel, Nikolay S. Strigul

## Abstract

Forest dynamics are highly dimensional phenomena that are not fully understood theoretically. Forest inventory datasets offer unprecedented opportunities to model these dynamics, but they are analytically challenging due to high dimensionality and sampling irregularities across years. We develop a data-intensive methodology for predicting forest stand dynamics using such datasets. Our methodology involves the following steps: 1) computing stand level characteristics from individual tree measurements, 2) reducing the characteristic dimensionality through analyses of their correlations, 3) parameterizing transition matrices for each uncorrelated dimension using Gibbs sampling, and 4) deriving predictions of forest developments at different timescales. Applying our methodology to a forest inventory database from Quebec, Canada, we discovered that four uncorrelated dimensions were required to describe the stand structure: the biomass, biodiversity, shade tolerance index and stand age. We were able to successfully estimate transition matrices for each of these dimensions. The model predicted substantial short-term increases in biomass and longer-term increases in the average age of trees, biodiversity, and shade intolerant species. Using highly dimensional and irregularly sampled forest inventory data, our original data-intensive methodology provides both descriptions of the short-term dynamics as well as predictions of forest development on a longer timescale. This method can be applied in other contexts such as conservation and silviculture, and can be delivered as an efficient tool for sustainable forest management.

## Software and data availability

The software to estimate transition matrices based on forest inventory was implemented by Jean Liénard in R version 2.15.1 (R Core Team, 2012) and is attached as a zip file to the submission.

The database studied in this paper is available upon request to the Quebec provincial forest inventory database (http://www.mffp.gouv.qc.ca/forets/inventaire/). Straightforward modifications of the software allows to use with the USDA Forest Inventory and Analysis program (http://www.fia.fs.fed.uss/).

## 1. Introduction

Forest ecosystems are complex adaptive systems with hierarchical structures resulting from self-organization in multiple dimensions simultaneously (Levin, 1999). The patch-mosaic concept was actively developed in the second half of the twentieth century after Watt (1947) suggested that ecological systems can be considered a collection of patches at different successional stages. Dynamical equilibria arise at the level of the mosaic of patches rather than at the level of one patch. The classic patch-mosaic methodology assumes that patch dynamics can be represented by changes in macroscopic variables characterizing the state of the patch as a function of time (Levin and Paine, 1974). Forest disturbances are traditionally associated with a loss of biomass; however, Markov chain models based only on biomass do not capture forest succession comprehensively (Strigul et al., 2012). This limitation motivates the need for alternative formulations that are able to consider several forest dimensions instead of only one.

Here we develop a novel statistical methodology for estimating transition probability matrices from forest inventory data and generalize classic patch-mosaic framework to multiple uncorrelated dimensions. In particular, we develop a landscape-scale patch-mosaic model of forest stand dynamics using a Markov chain framework, and validate the model using the Quebec provincial forest inventory data. The novelty of our modeling framework lies in the consideration of forest transitions within multiple dimensional space of macroscopic stand-level characteristics (biomass, average age of trees, biodiversity and shade tolerance index) that constitutes a generalization of the one-dimensional model of forest biomass transitions developed earlier (Strigul et al., 2012). Our framework is also substantially distinct from previous models of forest dynamics, where successional stages are ordinated using empirical observations on successional pathways (Curtis and McIntosh, 1951; Kessell and Potter, 1980; Logofet and Lesnaya, 2000).

The Quebec forest inventory (Perron et al., 2011) is one of the extensive forest inventories that have been established in North America, among others led by the Canadian provincial governments and the USDA Forest Inventories and Analysis program in the USA. These inventories provide a representative sample of vegetation across the landscape through a large number of permanent plots that are measured repeatedly. Although they were originally developed for estimating growth and yield, they were rapidly found to be extremely useful to studies in forest ecology, biogeography and landscape dynamics. Each permanent plot consists of individually marked trees that are periodically surveyed and remeasured. Each plot can be considered as a forest stand and then, theoretically, the forest inventories provide empirical data sufficient for parametrization and validations of patch-mosaic models (Strigul et al., 2012). However, practical development of the patch-mosaic forest models (i.e. their parametrization, validation and prediction) is challenging due to the underlying structure of the forest inventory datasets. These datasets are indeed collected at irregular time intervals that are not synchronized across the focal area, and data collection procedures including spatial plot design and tree measurement methods can be different at various survey times and conducted by different surveyors (Strigul et al., 2012).

Our objective in this study is to develop a data-intensive method predicting the dynamics of forest macroscopic characteristics. The idea of a data-intensive modeling approach is to develop and explore a quantitative theory using statistical modeling, in contrast with the hypothesis-driven theoretical approach in which selected mechanisms are used to design and constrain models. We focus here on the development of the modeling framework and illustrate the application of the framework to a large forest inventory dataset spanning 38 years of observations collected in Quebec. To overcome the issue of irregular samplings in time specific to forest inventory data, we develop a Gibbs sampling procedure for augmenting the data and infer the transition probabilities. Our particular use of Gibbs sampling (Pasanisi et al., 2012) has a substantial scientific novelty, as this is the first application of this statistical machinery to overcome the problems of irregularities in the forest inventory sample design. In this paper, we demonstrate the power of this statistical methodology in our application, and deliver it as ready-to-go tool for other applications by explaining every step, providing pseudocode, and original R code. We anticipate that this novel statistical methodology will be broadly used in forest inventory analysis as the issue of irregularities in inventories has previously been a substantial hindrance (e.g. Strigul et al., 2012).

We present in this paper the general methodology and demonstrate each of its steps on the Quebec dataset. In particular, we consider the dimensionality of stand characteristics in this dataset and present evidence that some characteristics are redundant. We apply the method to predict long-term dynamics of Quebec forests, as represented by a subset of macroscopic properties that best represent the variability in the data. We validate the model utilizing two different cross-validation schemes to split the original data, based on survey date (predicting later years using earlier years) and based on a random 2-folds partition of plots (comparing long-term predictions inferred from two independent subsets). We finally discuss the implications of this work, such as the effect of spatial and temporal variability, the independence of most forest variables, the effect of changing external drivers and of feedbacks.

## 2. Patch-mosaic modeling framework

The goal of this section is to introduce the modeling of patch-mosaic using Markov chains, which is generalized and employed to predict forest dynamics in the main text. The patch-mosaic concept assumes that the vegetation at the landscape level can be represented as a collection of isolated spatial units - patches - where patch development follows a general trajectory and is subject to disturbances (Watt, 1947; Levin and Paine, 1974). Patch-mosaic models are derived using the conservation law, which takes into account patch aging and other changes to macroscopic variables representing succession, growth of patches in space, and disturbances (Levin and Paine, 1974). The same general idea as well as mathematical derivations are broadly used in population dynamics to describe age- and size-structured population dynamics. Patch-mosaic models can be partial differential equations or discrete models depending on whether time and patch stages are assumed to be continuous or discrete. Classic continuous patch-mosaic models are based on the application of the conservation law to continuously evolving patches that can be destroyed with a certain probability, and can be represented by the advection equation (model developed by Levin and Paine, 1974, for fixed-size patches) or equivalently by the Lotka-McKendick-von Foerster model (Strigul et al., 2008). The continuous patch-mosaic models have been used in forest ecology to model the dynamics of individual canopy trees within the stand or forest gap dynamics (Kohyama et al., 2001; Kohyama, 2006).

In the case of patches changing in discrete time, the derivation of the conservation law leads to discrete-type patch-mosaic models. In particular, the advection-equation model (Levin and Paine, 1974) is essentially equivalent to several independently developed discrete models (Leslie, 1945; Feller, 1971; Van Wagner, 1978; Caswell, 2001). These models consider only large scale catastrophic disturbances (patch “death” process), destroying the patch, which then develops along the selected physiological axis until the next catastrophic disturbance Levin and Paine (1974). The stochastic model we are considering here employs a Markov chain framework (Waggoner and Stephens, 1970; Usher, 1979a; Facelli and Pickett, 1990; Logofet and Lesnaya, 2000; Caswell, 2001) that is capable of taking into account all possible disturbances.

In a Markov chains model, the next state of a forest stand depends only on the previous state, and the probabilities of going from one state into another are summarized in what is called a transition matrix, denoted *T*.

We summarize the distribution of states at time *t* as the row vector *X*_*t*_, with length equal to the number of discrete classes of patch state and with a sum equal to 1. We can predict *X*_*t*+Δ*t*_ by multiplying the transition matrix:

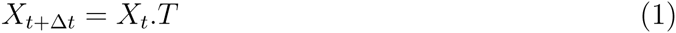

To project an arbitrary number *n* time steps into the future, one simply multiplies by *T*^*n*^ instead of *T*. The Perron-Frobenious Theorem guarantees the existence of the long-term equilibrium, which can be practically found as the normalized eigenvector corresponding to the first eigenvalue, or by iterative sequence of state vectors. In this paper we employ the iterative method as it allows to derive forest states at different time steps in the future, for example allowing to make predictions in 10, 20 or 30 years from now. To derive the long-term equilibrium we simply choose an *n* large enough to satisfy the condition:

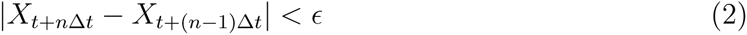

Three illustrative examples of simplified Markov chains are available in Appendix 1.

## 3. Materials and Methods

We address here the issue of constructing transition matrices from forest inventory data stemming from irregular sampling intervals and variable numbers of plots sampled in each year. We outline in the following the general concepts of the methodology along with practical guidelines using the inventory led by the provincial Ministry of Natural Resources and Wildlife in Quebec (Appendix 1). The key steps to use Gibbs sampling to estimate a transition matrix from irregular measurements are:

1. Compute stand level characteristics for each plot and for each survey year. Analyze the dimensionality of these characteristics using correlation and principal component analysis;
2. Construct temporal sequences of uncorrelated characteristics depending on forest survey dates. Use Gibbs sampling to infer the transition matrix. This algorithm consists of random initialization of missing values followed by iteration of parameter estimation and data augmentation:
  - Parameter estimation: Compute the transition matrix using the (augmented) sequences of plot transitions.
  - Data augmentation: Draw new sequences conditional on the new transition matrix.

The transition matrices for Quebec forests were obtained using this method with a three-year time step. Future and equilibrium landscape characteristics were predicted according to equations 1 and 2 (cf. Appendix 1).

### 3.1. Step 1: stand characteristics and dimensional analysis

This step consists of (a) the selection of a set of stand-level forest characteristics, (b) the dimensional analysis of these characteristics, (c) their decomposition into uncorrelated axes, and (d) the discretization of these uncorrelated axes.

Our modeling method can be applied for the prediction of any forest stand characteristic under the condition that it is computable from every single plot survey. The particular choice of the characteristics depends on available data and research objectives. A general guideline is that these characteristics should summarize data from individual trees into macroscopic indicators of stand structure, which can then be used to compare forests across different ecosystems. We consider six characteristics of Quebec forests according to the rationale presented in Strigul et al. (2012) and Lienard et al. (2014). These characteristics are computed based on trees with a diameter at breast height larger than 90mm (see Appendix 1.1 for more details about the Quebec forest inventory measuring protocol). We denote $ the set of species inside each plot and 𝕋 the set of trees inside each plot, and compute for each single plot survey the following characteristics:

- dry weight biomass, estimated from Jenkins et al. (2003), using the formula: 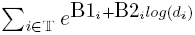 where B1 and B2 are species specific density constants, and *d* is the trunk diameter at breast height in cm. B1 and B2 have been derived from both US and Canadian studies, making it a suitable approximation for Quebec forests (Jenkins et al., 2003). The resulting aboveground biomass is expressed in 10^3^ kg/ha.
- basal area, computed as the sums of trunk diameters at breast height *d*: 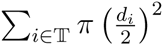. The basal area is expressed in m^2^/ha.
- intra-plot diversity (evenness), computed as the Gini-Simpson index (Hill, 2003), with Ω(*s*) referring to the number of trees with species *s* and Ω (𝕋) referring to the total number of trees inside each plot: 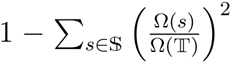. This provides an index in the 0-1 range describing the species heterogeneity at the stand level, with high values indicating a high heterogeneity.
- extra-plot diversity (species richness), computed as the number of species present in a plot: Ω($). In the Quebec dataset, this indicator ranges from 1 to 8 species, and is interpreted as another measure of diversity.
- shade tolerance index, a new metric introduced by Strigul and Florescu (2012) and Lienard et al. (2014) describing the shade tolerance rank of species 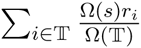. This index ranges from 0 to 1, with high values denoting forest stands composed of typically late successional species and low values denoting forest stands composed of typically early successional species in Quebec (Lienard et al., 2014).
- average age, computed as the average of tree ages *a* :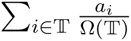. This commonly-used indicator approximates the stand age in the forest inventory analysis (see Strigul et al. 2012 for a discussion of this characteristic).

Statistical relations of these stand-level characteristics were analyzed using standard multivariate methods. First, we computed the Pearson correlation coefficients both in the whole dataset and in the dataset broken down in decades (to avoid biases due to their temporal autocorrelation). We then performed a principal component analysis (PCA) to examine (a) the number of components needed to explain most of the variance as well as (b) the projection of characteristics in the space defined by these components.

In general, it is possible for a multidimensional model to operate on the space of principal components. Such a model would (a) project the characteristics into the low-dimensional space given by the principal components, then (b) predict their dynamics in this new space, and finally (c) perform the inverse transformation to obtain predictions on the characteristics. In our application to the Quebec dataset, we discovered that four uncorrelated characteristics approximate well the principal component space (namely biomass, average age of trees, Gini-Simpson and shade tolerance indexes, *cf.* Results). Our model employs this approximation and is based on transition matrices of these forest characteristics. It substantially simplify interpretation of modeling predictions.

Prior to the computation of transition matrices in the Markov chain framework, it is necessary to discretize continuous variables into distinct states (Strigul et al., 2012). The general approach is to subdivide data into uniformly spaced states, with a precision that is small enough to capture the details of the distribution but large enough to be insensitive to statistical noise in the dataset. In addition, the computational effort needed to infer transition matrices is proportional to the square of the number of states, and available computational power may constitute a practical limitation to the number of states. In the Quebec dataset, the stand-level characteristics span different ranges (see Figs. 2 and 3 in Appendix), with the biomass distribution in particular showing a long tail for the highest values. In order to capture enough details of the distributions of the Quebec characteristics, we opted to remove plots in the long tail of the biomass (those with a biomass higher than 50,000 kg/ha, representing roughly 4% of the total dataset) and then subdivided the remaining plots into 25 biomass states. An alternative approach would be to merge the rarely occurring high-biomass states into the last state as was implemented in Strigul et al. (2012). We conducted a comparison of these two approaches and found no significant differences. For the other characteristics investigated (i.e. the internal diversity, shade tolerance index, and average age), we found that 10 states were enough to capture their distributions with sufficient detail.

### 3.2. Step 2: Gibbs sampling methodology

Inferring a Markov Chain model for characteristics computed with field data sampled at irregular intervals is a challenging problem. Indeed, the usual direct approach of establishing the *n*-year transition matrix by simply counting the number of times each state changes to another after *n* years can not be employed in most forest inventories, as successive measurements on the same plot are not made with constant time intervals. This irregularity in sampling results in states of the forest plots that are not observed, and can be modeled as missing data. Two classes of algorithms can be used to parameterize a transition matrix describing the dynamics of both observed and missing data: expectation-maximization (EM) and Monte Carlo Markov Chain (MCMC), of which Gibbs sampling is a specific implementation. Both classes of algorithms are iterative and can be used to find the transition matrix that best fits the observed data. EM algorithms consist of the iteration of two steps: in the expectation step the likelihood of transition matrices is explicitly computed given the distribution of the missing data inferred from the previous transition matrix estimate, and in the maximization step a new transition matrix maximizing this likelihood is chosen as the new estimate (Dempster et al., 1977). MCMC algorithms can be seen as the Bayesian counterpart of EM algorithms, as at each iteration a new transition matrix is stochastically drawn with the prior information of estimated missing data, and in turn new estimates for the missing data are stochastically drawn from the new transition matrix (Gelfand and Smith, 1990). EMs are deterministic algorithms, and as such they will always converge to the same transition matrix with the same starting conditions; conversely, MCMCs are stochastic and are not guaranteed to converge toward the same estimate with different random seeds. While both algorithms are arguably usable in our context, the ease of implementation and lower computational cost of MCMC algorithms led us to prefer them over EM (Deltour et al., 1999). We selected Gibbs sampling as a flexible MCMC implementation (Geman and Geman, 1984). We provide in the following a brief presentation of Gibbs sampling. Additional implementation details are in Appendix 1.2, and we refer to Robert and Casella (2004) for the general principles underlying MCMC algorithms and to Pasanisi et al. (2012) for an extended description of Gibbs sampling to infer transition probabilities in temporal sequences. In addition to the full explanation below, we also provide a pseudocode of the procedure (Box 1).

To apply Gibbs sampling for the estimation of the transition matrices, it is required to include plot characteristics in a set of temporal sequences. For each plot *p*, this is done by inserting each characteristic *s*_(*p,i*)_ measured in the *i*-th year at position *i* of a row vector *S*_*p*_ representing the temporal sequence of this plot. For example, if a plot *p* was sampled only at years 1 and 3 during a 5-year inventory, allowing for the computation of characteristics *s*_(*p,*1)_ and *s*_(*p,*3)_, then its sequence would be the row vector *S*_*p*_ = [*s*_(*p,*1)_, *•, s*_(*p,*3)_, •, •], where • denotes a missing value. The sequences are mostly composed of unknown values as only a fraction of the forest plots were surveyed each year. In the application to the Quebec dataset, a reduction of the size of these temporal sequences was performed (see Appendix 1.2 for a detailed description of this reduction and an illustrative example); however it is not a pre-requisite for the general application of Gibbs sampling. Let further *Y* be the matrix constructed using all the sequences *S*, with rows corresponding to successive measures of different plots and columns corresponding to different years. The preliminary step of Gibbs sampling consists of replacing the missing values • in *Y* at random, resulting in so-called augmented data *Z*^[0]^. Then, the two following steps are iterated a fixed number of times *H*, with *h* the index of the current iteration:

1. 1. in the **parameter estimation** step, we draw a new transition matrix Φ ^[h]^ conditional on the augmented data *Z*^*h*–1^, using for every row *i*:

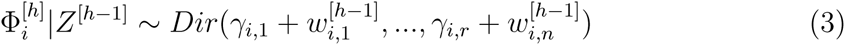

with *Dir* is the Dirichlet distribution, γ are biasing factors set here uniformly to 1 as we include no prior knowledge on the shape of the transition matrix (Pasanisi et al., 2012). *w*_*i,j*_ are the sufficient statistics reflecting the transitions in the augmented data *Z*^[h–1]^, formally defined as

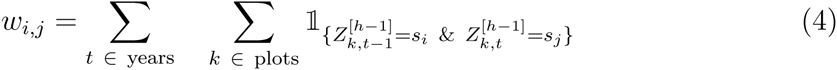

with 1_{*Y*_*k,t−*1_=*s*_*i*_ & *Y*_*k,t*_=*s*_*j*_}_ the count of sequences elements in the state *s*_*i*_ at time *t* − 1 and in the state *s*_*j*_ at time *t*.
2. in the **data augmentation** step, we draw new values the missing states, based to the probabilities of the transition matrix Φ^[h]^. The probabilities ℙ used to augment the data *Z*^[h]^ are derived from their values in the previous iteration (*Z*^[h–1]^) as well as their values in the current iteration but in an earlier year 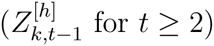: 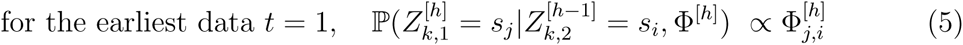

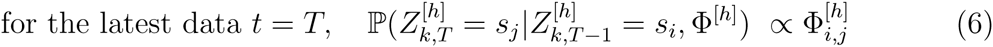

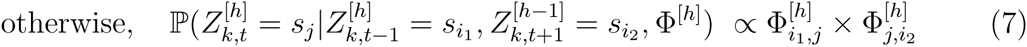

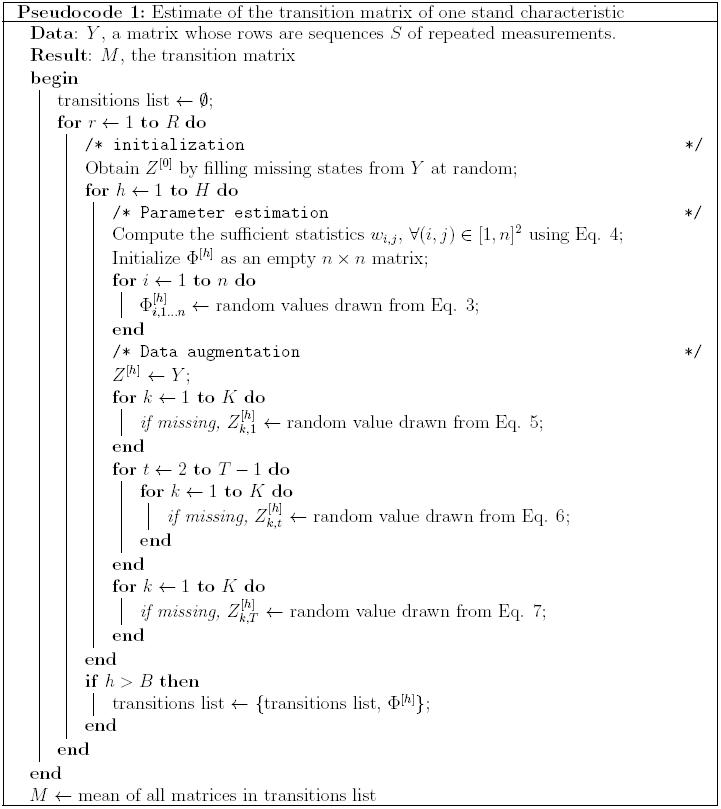

As Gibbs sampling is initialized by completing the missing values at random, the first iterations will likely result in transition matrices far away from the optimal. The usual workaround is to ignore the first *B* transition matrices corresponding to so-called “burn-in” period, leaving only *H* – *B* matrices. Furthermore, as Gibbs sampling relies on a stochastic exploration of the search space, a good practice to ensure that Gibbs sampling converged to the optimal transition matrix is to run the whole algorithm *R* times. There are no general guidelines for setting the *H*, *B* and *R* parameters (Robert and Casella, 2004). We empirically settled with *H* = 1000, *B* = 100 and *R* = 50 in order to ensure that the transition matrices were reproducible for the Quebec dataset, leading to *R* × *H* = 50000 iterations of parameter estimation and data augmentation steps and resulting in *R* × (*H − B*) = 45000 transition matrices. This process was repeated independently for each plot characteristic. The algorithm was implemented in R version 2.15.1 (R Core Team, 2012) and took a total runtime of 4 days on a 1.2 Ghz single-core CPU to compute the transition matrices for all 4 characteristics studied here.

## 4. Results

### 4.1. Multivariate analysis of stand characteristics

The correlation analysis performed on the Quebec forest inventory (Perron et al., 2011, Appendix 1.1) revealed that biomass and basal area were highly correlated (*r* = 0.96), as well as the external and internal diversity indices (*r* = 0.90, see Appendix 1.3 for the other coefficients). These correlations are further preserved when the correlation analysis is done separately on each decade, from the 1970s until the 2000s (*cf*. tables in Appendix 1.3), confirming the presence of time-independent strong correlations between these two pairs of characteristics.

A PCA applied to the dataset further confirmed that the biomass and basal area on one hand, as well as the external and internal diversity on the other hand, have nearly identical vectors in the principal components space (*cf*. Appendix 1.4). Furthermore, this analysis showed that 4 principal components are required to adequately explain variance in the data; using 3 components accounts for only 87 % of the variance, while 4 components explain up to 98 % of the variance. The PCA revealed that biomass, the internal diversity index, the shade tolerance index, and the average age are close approximations of the different principal components and explain most of the variance. Therefore, these variables have been employed in the following analysis.

### 4.2. Interpretation of the transition matrices

We present here in detail the transition matrix for biomass with a 3-year time interval, shown in Fig. 1 (the other characteristics are to be found in Appendices, in Figs. 7 and 8). In this matrix, each value at row *i* and column *j* corresponds to the probability of transition from state *i* into state *j* after 3 years. By definition, rows sum to 100%. This transition matrix, as with the others in Appendix, is dominated by its diagonal elements, which is expected because few plots show large changes in a given 3-year period. The values below the diagonal correspond to transitions to a lower state (hence, they can be interpreted as the probabilities of disturbance), while values above the diagonal correspond to transitions to a higher state (i.e., growth). The transitions in the first column of the matrix correspond to major disturbances, where the stand transitions to a very low biomass condition. As the probabilities above the diagonal are larger than below the diagonal, the overall 3-year prediction is of an increase in biomass. This matrix also shows that plots with a biomass larger than 40,000 kg/ha have a roughly uniform 10% probability of ending with a biomass of less than 20 000 kg/ha 3 years later, which is interpreted as the probability of high-biomass stand to go through a moderate to high disturbance.

**Fig. 1.**
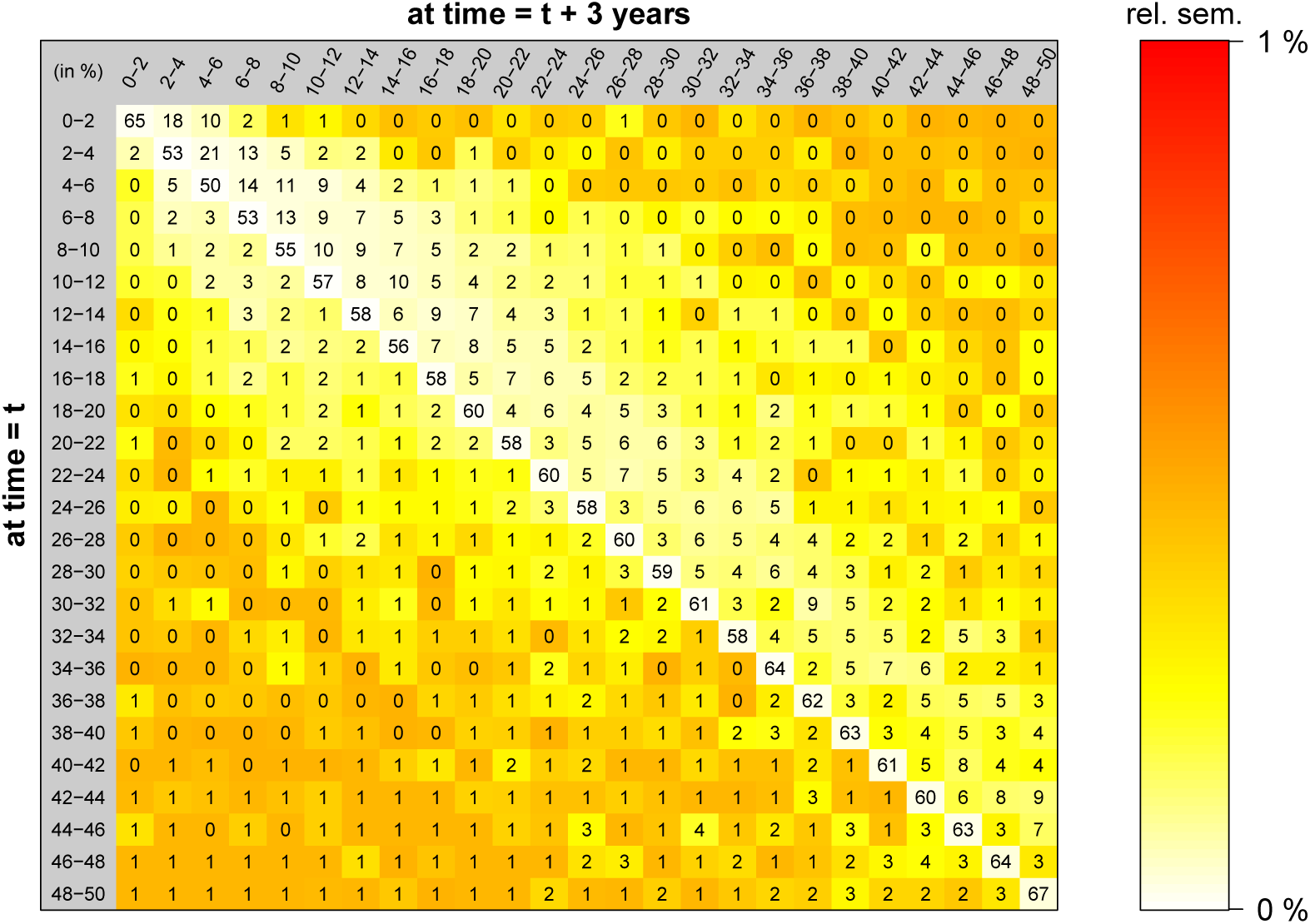
3-year transition matrix for the biomass. The states are the biomass ranges in 10^3^ kg/ha, spanning from 0 − 2 to 48 − 50 10^3^ kg/ha, and represented here on the left and on top of the matrix. The values *M*(*i, j*) inside the matrix correspond to the rounded probability of transition from state *i* to state *j*. The color represents the relative standard error of the mean and indicates the robustness of the stochastic search, as explained in Section 4.3. Lighter colors thus indicate a better confidence in the transition value; all relative standard errors of the mean (RSEM) are below 1%, corresponding to a very high confidence, and furthermore the smallest errors are found for the higher transition probabilities close to the diagonal.

### 4.3. Model validation

Two main types of error should be considered when designing a model with a parameter search based on real data. The first error relates to the robustness and efficiency of the estimation of the optimal transition matrix, which was performed with Gibbs sampling in our case. The second type of error encompasses more broadly the capacity of the chosen theoretical framework to predict the system beyond the range of the dataset. In our case, the theoretical framework we relied on is patch-mosaic concept, implemented with the Markov chain machinery, to describe the dynamics of our four characteristics.

To estimate the errors of the parameter search, we used the *R*(*H* − *B*) transition matrices to compute for each transition the standard error of the mean (SEM) and the relative standard error of the mean (RSEM, defined as the ratio of the SEM over the transition probability, and expressed as a percentage). The SEM were below 1% throughout the matrices, with the highest errors occurring for very low transition probabilities (i.e., far from the diagonals). Furthermore, the RSEM were very low, and particularly so for the transitions with the highest probability (Fig. 1 in main text as well as Figs. 7 and 8 in Appendix). We finally computed the SEM in the long-term predicted equilibriums and found values below 0.01%, strengthening the conclusion that negligible errors are to be attributed to the stochastic fit procedure.

An independent dataset would be most suited to estimate the more general errors in the ability of a Markov-Chain model to predict future forest characteristics. As there is no such dataset available, we performed two cross-validations of our methodology by splitting this dataset in two different ways. In the first, we ran the Gibbs sampler with only the first 18 years of records (from 1970 to 1988). We then used the model to predict forest state for the period corresponding to the second half of the dataset (i.e., 1989 to 2007), and we compared the predicted dynamics with the aggregated distribution of the second half of the dataset (Fig. 2). Overall, the predictions were highly accurate, with *R*^2^ between observation and prediction ranging from 0.8 to 0.95, indicating that the second half of the dataset is predictable with a Markov chain model based solely on the first half. In the second validation, we randomly split the data into two sets, regardless of year. We then computed the transition matrix and corresponding equilibrium conditions for each half (Fig. 9 in Appendix). Here again, the predictions match closely with values of *R*^2^ higher than 0.98 for the internal diversity, shade tolerance index and average age. The *R*^2^ was near 0.6 for the biomass, indicating that this variable is more sensitive to small changes than the others; however the difference in predictions were small, typically around 1% for each biomass state. This second validation overall showed that the data contained in the inventory is redundant, and that half of it is enough to provide highly accurate long-term estimates for the internal diversity, shade tolerance index and average age. Considering only half of the data at random would likely result in errors of around 1% in the long-term estimates of the biomass.

**Fig. 2.**
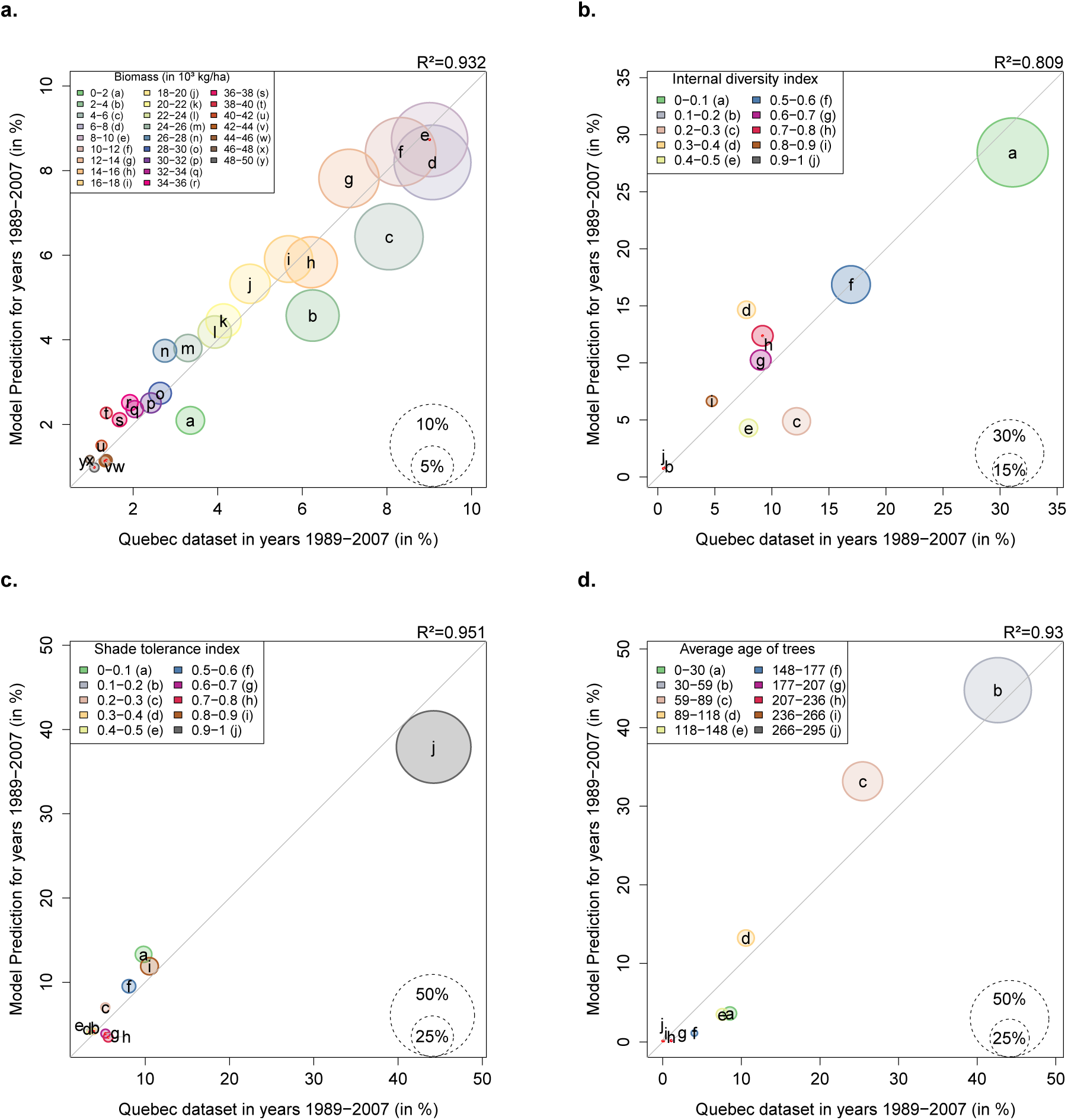
Results of model validation, showing the second half of the dataset *vs* the model prediction for the classes of each characteristic (distribution in %). For each class, the circle size denotes the number of stands belonging to it in the real dataset. The *R*^2^ measure is indicated on the top right of each plot. The model used to make the prediction was computed using only the first half of the dataset, corresponding to years 1970 to 1988 (see Materials & Methods for details).

### 4.4. Predictions of temporal dynamics and long-term equilibrium

We applied the inferred transition matrix to predict the state of forest in 2010s, 2020s and 2030s based on their distribution in 2000s. We also predicted the long-term dynamics of the forest stands, by computing the equilibrium states of the transition matrices. Overall, the predictions showed an increase in biomass and stand age (Fig. 3 e and h), along with a slight increase in biodiversity (Fig. 3 f) and a slight decrease of the prevalence of late successional species accompanied by a slight increase of early successional species (Fig. 3 g). These predictions are obvious for the biomass and average age of trees by looking at their distributions in the existing dataset (Fig. 3 a and d), while they are less clearly seen when looking at the average distributions of the biodiversity and shade tolerance index (Fig. 3 b and c).

**Fig. 3.**
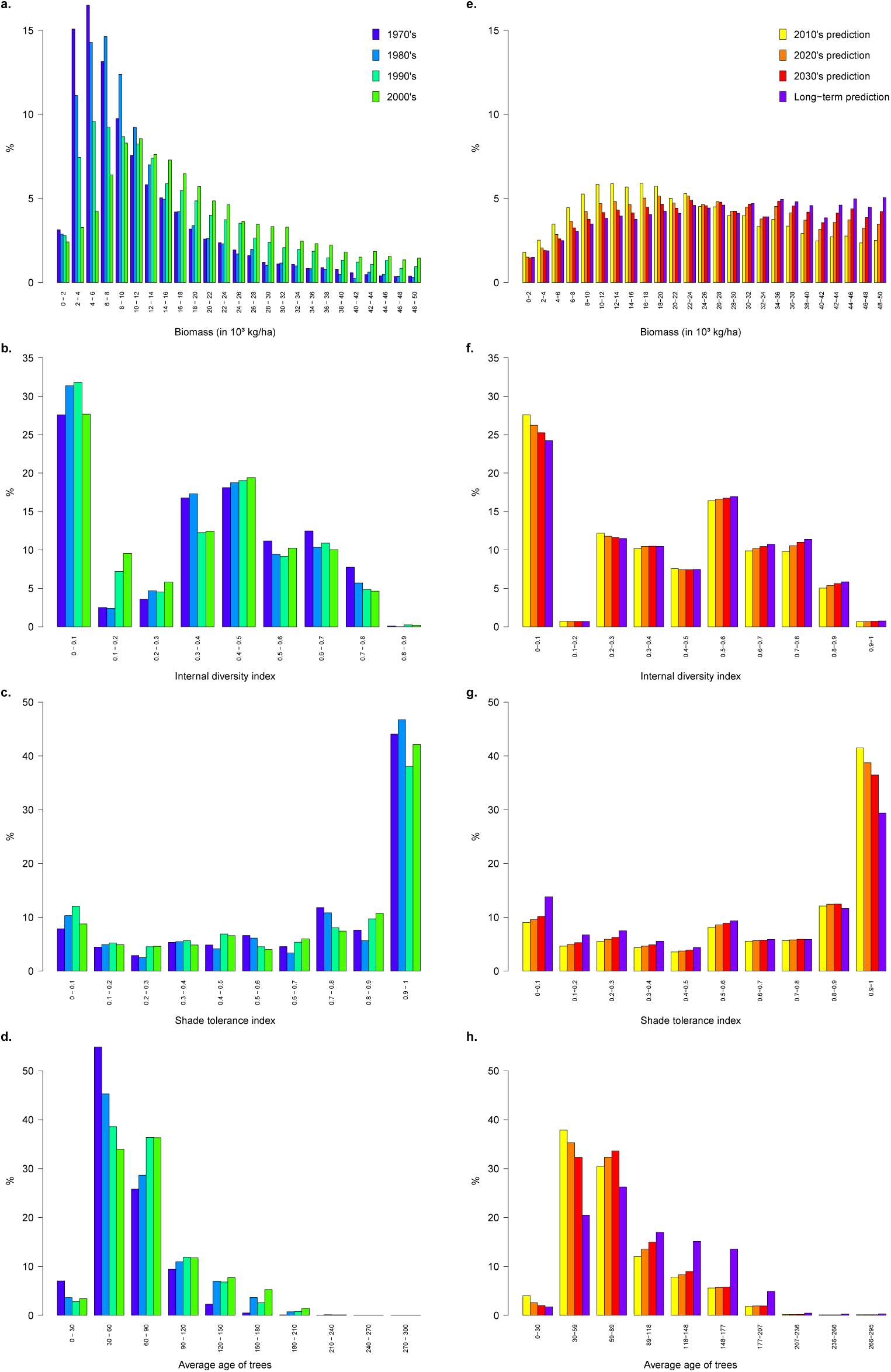
Current distribution of relevant characteristics from the database, along with the long-term predictions of our models.

These long-term predictions were reached at different timescales depending on the characteristics. For biomass, equilibrium was reached by approximately year 2030, but the other characteristics, and in particular the average age of trees in plots, showed much slower dynamics to reach their equilibria (Fig. 3 e to h). The model predicted average relative changes of +38.9% and +14.2% by the 2030s for biomass and stand age, and +44.0% and +37.9% by the time they reach their long-term equilibrium state. Relative changes for the Gini-Simpson diversity index were +5.2% by the 2030s and +7.1% in the long term, and early successional species will become slightly more abundant with a change of -4.7% of the shade tolerance index by the 2030s and -13.6% in the long term.

## 5. Discussion

We developed a data-intensive approach to multiple-dimensional modeling of forest dynamics. The modeling steps include 1) dimensional analysis of forest inventory data, 2) extraction of non-correlated dimensions, and 3) the application of stochastic optimization to compute probability transition matrices for each dimension. We applied this approach to the Quebec forest inventory dataset and validated the model using two independent subsets of data. Our study demonstrates that there exist at least four uncorrelated dimensions in Quebec forests: the biomass, biodiversity, shade tolerance index and averaged age of trees. The most pronounced changes predicted for Quebec forests are increases in biomass and stand age. Our model also predicted smaller increases in biodiversity in the prevalence of early successional species. Our results demonstrate the utility of this methodology in predicting long-term forest dynamics given highly dimensional, irregularly sampled data; the model was computationally efficient and validation procedures demonstrated its ability to make short and long-term predictions. Therefore, the framework will be useful both in applied contexts (e.g., conservation, silviculture) as well as in developing our conceptual understanding of how forested ecosystems are organized through dimensional analysis of forest characteristics under the current disturbance regime.

### 5.1. Contribution to Markov chain forest modeling framework

Markov chain models have a rich history of application in ecology, and, in particular, in forest modeling (Facelli and Pickett, 1990; Caswell, 2001). This modeling framework has been employed to describe forest transitions at different scales with various focal variables, for example, succession models defined on the species and forest type level (Usher, 1969, 1981; Waggoner and Stephens, 1970; Horn, 1974; Logofet and Lesnaya, 2000; Korotkov et al., 2001), gap mosaic transition models (Acevedo et al., 1996, 2001) and biomass transition models (Strigul et al., 2012). Markov chain successional models (Usher, 1969, 1981, 1979b; Facelli and Pickett, 1990) are able to predict changes in species abundance, but require a comprehensive knowledge of successional sequence of species replacement and transition probabilities between different successional stages. The empirically-based Markov chain forest succession model, which operates at the species level and assumes that the underlying Markov chain is stationary, requires only substantially large observations to estimate transition probabilities (Waggoner and Stephens, 1970; Stephens and Waggoner, 1980). On the contrary, the mechanistic Markov chain modeling approach developed by Horn (1974, 1981) employs shade tolerance and gap dynamics to predict species replacement in the forest canopy given the species composition in the understory. However, this approach requires a detailed survey of the understory vegetation that is not commonly available in forest inventories. Also, gap dynamics individual-based models can be coupled with Markov chain models for scaling of gap dynamics to patch level (Acevedo et al., 1996, 2001). These transitional models have been demonstrated to be useful and relevant tools in forest prognosis, however their practical applications are often limited. The Bayesian methodology proposed in this study allows to extend the scope of transition matrices by allowing their computation directly from forest inventory data, with corresponding modifications of the R code provided as a supplementary material. The proposed methodology of matrix estimation could be employed to test the validity of the Markov chain homogeneity assumption.

### 5.2. A data-intensive approach to understand forest dynamics

Modeling complex adaptive systems such as forest ecosystems requires capturing the dynamics of biological units at multiple scales and in multiple dimensions (Levin, 1998, 2003). Ideally, a mechanistic model based on the physiological processes and interactions of individual organisms should simulate the observed forest structure and predict forest dynamics over different time horizons and environmental variables. However, such individual-based modeling is very challenging as interactions between individual organisms within the forest stand result in new properties at the stand level, where essential mechanisms, spatial dimensions of variables, and functional relationships between variables are largely unknown. Given these unknowns, a data-intensive approach can be useful for gaining insight into ecosystem dynamics provided that sufficient amounts of relevant data are available (Kelling et al., 2009; Michener and Jones, 2012). In particular, the matrices we estimated (see Fig. 1 in main text and Figs. 7 and 8 in Appendices) incorporate all forest changes related to different magnitude disturbances. This opens a possibility of the future investigation of how particular disturbances are reflected in the forest macroscopic characteristics and can lead to a logical extension of classic models that take into account only major disturbances, in particular, birth-and-disaster Markov chains (Feller, 1971), forest fire models (Van Wagner, 1978), and advection-reaction equations for patch dynamics (Levin and Paine, 1974).

A potential limitation to a mechanistic interpretation of the transition matrices arises from the Markovian assumption that the transition toward the next state depends solely on the current state. If this assumption is not valid, it could bias these models. This assumption warrants further attention as it has not been yet comprehensively evaluated in forest modeling.

Integral Projection Model (IPM) is another modeling framework that could be used in place of Markov chains (Easterling et al., 2000; Caswell, 2001). In IPM, continuous kernel functions are used instead of discrete transition probabilities. While IPM are by design suited to handle well data irregularly distributed across the states, they do not address explicitly the issue of sampling irregularities in time. The data augmentation approach developed here can however be transposed to parameterizing IPM as well. Markov chains are preferable in our application because they are not restricted by the choice of IPM kernels. Indeed, biomass and stand age transitions can be decomposed into several kernels using commonly accepted assumptions of growth and disturbances, however there is no obvious way to choose kernels for biodiversity and shade tolerance index as their dynamics can not be understood in terms of a monotonic progression toward high values (Lienard et al., 2014).

The application of MCMC procedures allows to compute transition matrices for datasets with irregular sampling intervals and sample sizes. While Gibbs sampling has been introduced 30 years ago (Geman and Geman, 1984), its application to handle missing data in ecology has been mostly limited to stochastic patch occupancy models with a low number of free parameters (5-6) and either artificially simulated data or relatively restricted datasets (e.g. 72-228 resampled locations in ter Braak and Etienne, 2003; Harrison et al., 2011; Risk et al., 2011). From the technical point of view, our application of MCMC differs by taking advantage of the absolute time independence of Markov chains (allowing us to align subsequences starting with a known observation, see Methods and Appendix 1.1). This makes the use of MCMC possible in a data-intensive context, in which both the number of free parameters (600 for the biomass matrix, 90 for each of the biodiversity, shade tolerance and stand age matrices) and the number of samples (32,552) constitute increases of several orders of magnitude. Similar irregularity problems are quite common in ecological datasets, and the presented approach may have numerous applications beyond the statistical analysis of forest inventories. This methodology can also be applied to other datasets, even with regular samplings, and the same methodology can be applied to deduce transitions with a finer temporal scale.

In this study we have analyzed the Quebec forest inventories without explicitly taking into account the geographical location of plots, as well as the environmental and climatic variables. We have obtained transition matrices covering temperate to boreal forests, with a disturbance regime varying from canopy gaps to disastrous fires. We have repeated the developed approach after subdividing the Quebec dataset into the major ecological domains and have not observed substantial differences between the resulting transition matrices and the general matrices presented in this study (Liénard et al. unpublished data). In addition to this, the biomass transition matrices computed for the Lake States in the US (see Strigul et al. (2012) Tables 2 and 3) and the shade tolerance index transition matrices computed in northeastern parts of the US (Lienard et al., 2014) are quite similar to the ones presented in this study. It is quite amazing in fact that we could represent the dynamics of stand level characteristics given the neglection of geography. We hypothesize that the forest stand dynamics as well as disturbance regimes have substantial similarities across a large number of boreal and temperate forest types, and this will be specifically addressed in our future studies. We believe that the ability to make broad predictions on the forest stand dynamics without going into the fine details of geography is one of the major strengths of our approach.

The patch-mosaic framework has been already extensively employed in forest modeling (Kohyama et al., 2001; Kohyama, 2006; Scherstjanoi et al., 2013). Our approach has substantial similarities with the previous studies using the same scientific background (see Appendix 1), however there are distinctions related to the definitions of the forest patch. The forest stand or (forest patch) in this work represent a unit of forest which is large enough to be a community of trees, where individual tree gap dynamics is averaged, but at the same time small enough to be a subject to intermediate and large scale disturbances (Strigul et al., 2012)[p.72]. This definition results in an estimate of about 0.5-1 ha, which allows to use forest inventory permanent plots directly as an approximate forest stand representations (the size of the standard Quebec forest inventory plot is about 625 m^2^ and the USDA FIA plot is 675 m^2^). In other application of patch-mosaic concept to forest dynamics the patches (stands) are often defined differently. The size of patches varies from the size of large individual trees (in this case the patch dynamics is essentially equivalent to the gap dynamics Kohyama et al., 2001; Kohyama, 2006; Moorcroft et al., 2001), through patches similar to employed in our study (Acevedo et al., 2001) to the much large forest patches representing many hectares of forest (Boychuk et al., 1997). The difference in definitions of the patch essentially reflects the different applications and questions that can be addressed with particular models (see Strigul et al. (2012)[p.71] for an additional discussion).

### 5.3. Predictions for forest dynamics in Quebec

Our model made several notable predictions about future forest dynamics in Quebec. The most pronounced predicted changes are substantial short-term increase in biomass and a longer-term increase in average age of trees (Fig. 3). The increase in biomass is intuitively consistent with the increase in stand age, and both demonstrate a progression toward more mature stands. This progression is to be sustained throughout the next 20 years and beyond (Fig. 3), thus meaning that the unmanaged forests sampled in the inventory are currently far from their equilibrium state. The model also predicted smaller changes in biodiversity and the shade tolerance index. To understand stand maturation occurring with the small increase in the prevalence of early successional species, we must recall that neither biomass nor stand age are significantly correlated with shade tolerance index in the dataset (e.g., *r* = –0.02 with 95% confidence interval [-0.03,-0.01] for biomass and shade tolerance, see Fig. 5 in Appendix). Thus, it is unsurprising that the predictions are not correlated. Further, the predicted changes happen with different temporal dynamics and have different magnitudes, and have probably distinct mechanisms. In particular, while biomass and stand age are affected by both individual tree growth (leading to an increase) and disturbances (leading to a decrease), the shade tolerance index is affected only by disturbances. On the one hand, small disturbances (e.g., individual tree mortality) will typically promote the recruitment of late successional species into the canopy through gap dynamics. On the other hand, intermediate and large-scale disturbances will facilitate early successional species via the development of large canopy openings (e.g. Taylor and Chen, 2011). Thus, increase of intermediate and large-scale disturbances may promote early successional species, while the overall increase in biomass and stand age would result largely from individual tree growth. Our work thus suggests that Quebec forests are not progressing toward higher shade tolerance states despite their continuous biomass and stand age growth. This result echoes recent studies which showed that shade tolerance is not the sole driver for forest succession in Canadian central forests (Taylor and Chen, 2011; Chen and Taylor, 2012).

The accurate prediction of the second half of the dataset obtained using only the first half of the dataset demonstrate that the natural disturbance regime in the forest plots sampled in the Quebec inventory did not change substantially over the last 30 years. In the context of global warming, this could mean either that (a) there is no substantial consequence yet on the macroscopic dynamics of Quebec forests or that (b) the climatic change consequences were already present in the first half of the dataset or that (c) our analysis is not fine enough to catch the signal of the recent climate change (in particular moving climatic boundaries, cf. McKenney et al., 2007, are not taken into account as the approach developed here is not spatially explicit). In all cases, the inclusion in the transition matrices of future disturbances induced by climatic change (e.g. the increase of forest fire reviewed in Flannigan et al., 2009) could be a promising follow-up of our work by providing quantitative insights on the consequences of global warming on forests. The study of changes in disturbances was not the focus of the current study, and we have to be careful in generalizing conclusions about global warming as the gradual non-stationary disturbance regimes might take from 50 to 100 years to show significant departures (Loudermilk et al., 2013; Rhemtulla et al., 2009; Thompson et al., 2011).

The multidimensional nature of forest stands creates substantial challenges for modeling. Our study demonstrates that at least four dimensions are uncorrelated in the Quebec dataset, and that stand characteristics cannot be collapsed around one variable. The data intensive model could be based on uncorrelated principal component axes. However, such a model would not lend itself to a simple mechanistic interpretations in terms of macroscopic forest characteristics. Therefore, we have developed the model using the mostly uncorrelated stand characteristics: biomass, biodiversity, shade tolerance index, and average age of trees. In this model, the small correlations between these characteristics (Figure 3 in Appendix) will propagate to the model predictions, potentially resulting in slightly correlated predictions, in contrast with a model developed on the principal components. However, this choice of dimensional variables has the decisive advantage of allowing for the meaningful interpretation of the transition matrices and predictions.

The constructed transition matrices have predictive power, as demonstrated in Section 4.3. However, the universality of the predictions is intrinsically dependent on the representativity of the dataset, and a bias in data collection will be reported in the predictions. For this particular dataset for instance, we observed (and also predicted) increasing biomass, diversity, age and slight decreasing shade tolerance over time. However we should not expect forest stands affected by additional silvicultural operations, such as logging, to follow the trajectory recorded in the Quebec dataset. Thus, predictions made with this dataset should not be extended to them.

## Acknowledgements

This work was partially supported by a grant from the Simons Foundation (*#*283770 to N.S.) and a Washington State University New Faculty SEED grant. D.G. also acknowledges the financial support from a Strategic Grant of NSERC. We thank Matthew Talluto for interesting discussions and help with editing the manuscript.

